# Evolutionary dynamics in non-Markovian models of microbial populations

**DOI:** 10.1101/2022.06.17.496620

**Authors:** Farshid Jafarpour, Ethan Levien, Ariel Amir

**Affiliations:** Institute for Theoretical Physics, Utrecht University, Princetonplein 5, 3584 CC Utrecht, The Netherlands; Mathematics Department, Dartmouth College, Hanover, New Hampshire 03755, USA; John A. Paulson, School of Engineering and Applied Sciences, Harvard University, Cambridge, Massachusetts 02138, USA

## Abstract

In the past decade, great strides have been made to quantify the dynamics of single-cell growth and division in microbes. In order to make sense of the evolutionary history of these organisms, we must understand how features of single-cell growth and division influence evolutionary dynamics. This requires us to connect processes on the single-cell scale to population dynamics. Here, we consider a model of microbial growth in finite populations which explicitly incorporates the single-cell dynamics. We study the behavior of a mutant population in such a model and ask: can the evolutionary dynamics be coarse-grained so that the forces of natural selection and genetic drift can be expressed in terms of the long-term fitness? We show that it is in fact not possible, as there is no way to define a single fitness parameter (or reproductive rate) that defines the fate of an organism even in a constant environment. This is due to fluctuations in the population averaged division rate. As a result, various details of the single-cell dynamics affect the fate of a new mutant independently from how they affect the long-term growth rate of the mutant population. In particular, we show that in the case of neutral mutations, variability in generation times increases the rate of genetic drift, and in the case of beneficial mutations, variability decreases its fixation probability. Furthermore, we explain the source of the persistent division rate fluctuations and provide analytic solutions for the fixation probability as a multi-species generalization of the Euler-Lotka equation.

## I. INTRODUCTION

In recent decades, advances in single-cell technology and single-molecule biophysics have shed new light on the molecular mechanisms and dynamics underlying single-cell growth and division [1–3]. In some instances, aspects of cell growth are preserved across all domains of life, such as the adder mechanisms for cell-size regulation, which appears in bacteria, archaea and eukaryotes [4]. In other cases, even closely related organisms have found distinct ways to achieve the same objective, such as the regulation of mating type genes in yeast [5]. These observations have led to questions about how evolution has shaped single-cell physiology and vice-versa. While phylogenetic methods can, to some extent, help us understand the evolutionary history of physiological mechanisms, their applications are limited to cases where we have accesses to a clear mapping between the physiological mechanisms of interest and their associated genotypes. In order to understand cellular physiology from an evolution perspective, we must therefore develop a mechanistic understanding of how evolution acts on properties of single cells.

A standard model used in theoretical population genetics is the continuous time Moran process. In this model, cells of genotype *i* divide at a constant rate *ρ*_*i*_ (the fitness). In order to keep the population size fixed, every time a cell divides, a cell is randomly selected from the entire population (including the two newborn cells) to be expelled; see Figure 1 (A). The Moran process can be thought of as a mathematical idealization of a turbidostat culture – in such cultures the rate at which fresh media is continuously pumped into a vial is modulated in order to keep the optical density (and hence the number of cells) fixed. Since real organisms do not divide at a constant rate, *ρ* is typically thought of as a proxy for the viability of a genotype, rather than the literal division rate of a cell. Suppose that two genotypes are present in the population with fitnesses *ρ*_*r*_ (the resident) and *ρ*_*m*_ (the mutant). It can be shown that for sufficiently large populations, the fraction of mutant cells, denoted, *ϕ*, will obey the stochastic differential equation (SDE)

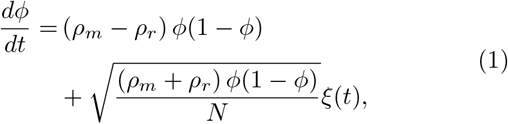

where the noise term should be interpreted in the Itô sense. It is from this SDE that all the fundamental results concerning the fate of rare mutations in an evolving population can be derived.

**FIG. 1.**
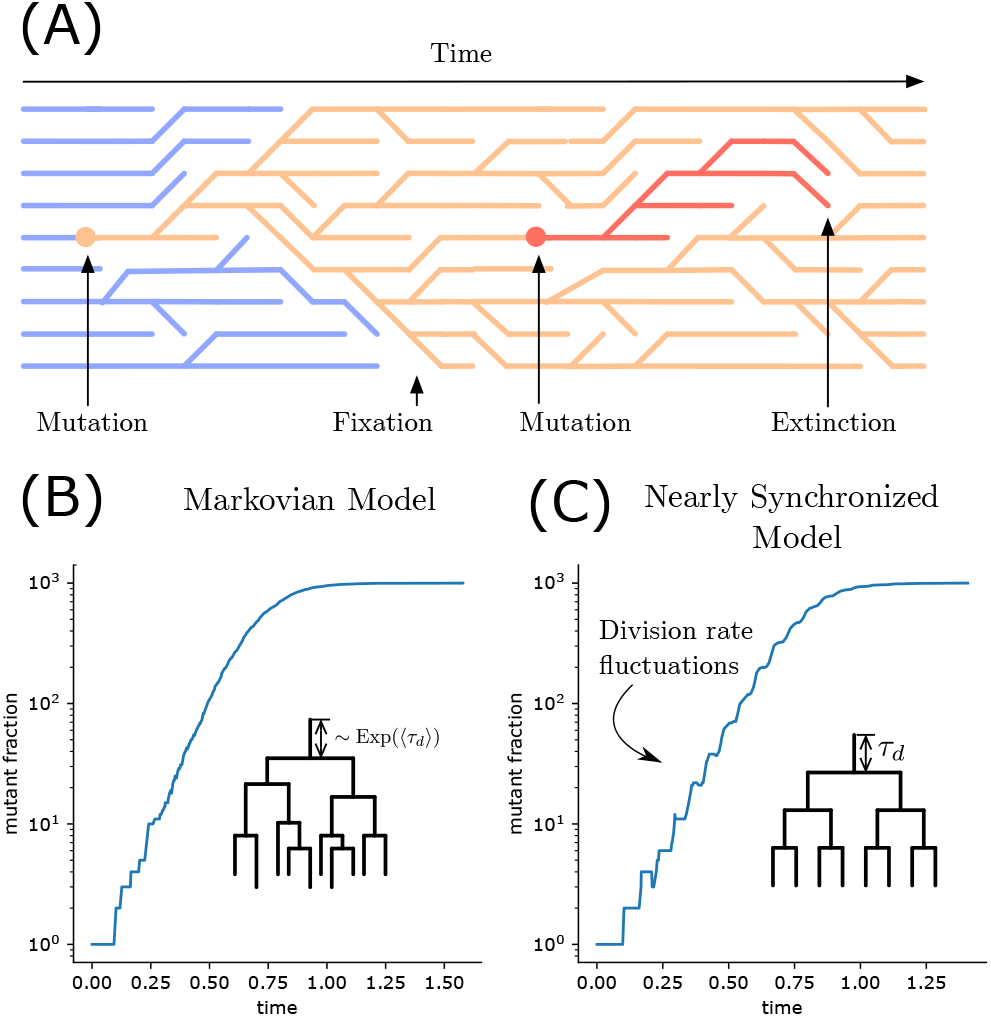
Panel (A) shows a schematics of the Moran process. After each division, a cell is expelled from the population to keep the total population size constant. Evolution proceeds by mutations that eventually either fix or go extinct. Panels (B) and (C) show trajectories of an initial mutant population which goes on to establish in the culture within two different models. The null model (B) is Markovian division where the probability of division for each cell is constant regardless of its size or age, corresponding to exponentially distributed generation times *τ*_*d*_. In this model, the rate of divisions of cells in the population quickly reaches a constant value. In a more realistic model of growth and division, panel (C), cell divisions remain nearly synchronized due to the narrow distribution of generation times *τ*_*d*_.

In the neutral case (*ρ*_*r*_ = *ρ*_*m*_ = *ρ*), the frequencies have mean ⟨*ϕ*⟩ = *ϕ*(0), while the variance in *ϕ* will evolve as

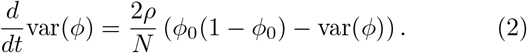

Therefore, the 2*ρ/N* factor sets the time-scale for the frequency of neutral mutations to change by a significant amount. We will derive Eq. (2) more generally in Appendix C1. In the case of a beneficial mutation (*ρ*_*m*_ *> ρ*_*r*_), when *ϕ >* 1*/Ns* (where *s* = 1 − *ρ*_*r*_*/ρ*_*m*_ is the *selection coefficient*), Eq. (1) is well approximated by a logistic growth [6]. However, since the mutant population initially consists of one cell, we are interest in the chance that the new mutant will eventually overtake the resident, or the *fixation probability*. A central result in population genetics states that in a large population the fixation probability is given by [7, 8]

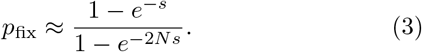

When *s* and *sN* are both sufficiently small, we obtain the simple relation *p*_fix_ ∼ *s*. Importantly, both the rate of genetic drift and the the probability for a mutant to fix depend only on the long-term growth rates of the resident and mutant populations, and therefore, one might conclude that the single-cell dynamics are relevant only insofar as they effect the long-term growth rate.

In order to develop a predictive theory of microbial evolution, we need to link the relevant parameters *ρ*_*r*_ and *ρ*_*m*_ to properties of single-cells. The problem of understanding how single-cell dynamics map to population fitness dates back to the work of Euler and Lotka [9]. By extending the work of these authors, recent studies have developed an understanding of how variation in traits such as cell-size, cell growth rates, and generation times combine to shape the long term growth of a population [10–19]. These studies have shown that under conditions of constant exponential growth, the distribution of ages in a population converges to a steady-state which determines the growth rate of a population. Moreover, the long-term fitness in the context of the Moran process (defined as the division rate, *ρ*) can be inferred from the distribution of generation times *ψ*(*τ*) measured among newborn cells using the Euler-Lotka equation:

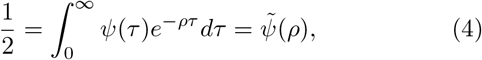

where 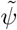 is the Laplace transform of *ψ*. This relation has proven in both experimental and theoretical studies to be a powerful tool for probing the effects of single-cell dynamics on growth. For example, it has been used to elucidate the effects of cell-size regulation in bacteria on population growth [12].

Equation (4), when combined with Eq. (1), appears to provide a link between the single-cell dynamics and the evolutionary dynamics; specifically, Eq. (4) can be used to obtain the long-term growth rates of each species which can then be used to find the rate of genetic drift and the fixation probability via Eq. (1). However, this reasoning ignores a crucial subtlety of this problem: Eq. (1) is derived under the assumption that there exists a constant division rate, *ρ*, which characterizes the fitness of a genotype in a finite culture. This assumption is valid in situations where *ϕ ≫* 1*/*(*Ns*), so that demographic fluctuations can be neglected. For rare mutants with small selections coefficients, or for neutral mutants, the division rate will fluctuate as a result of fluctuations in the distribution of cell ages. In this paper, we seek to understand the extent to which these fluctuations influence populations genetics.

Our focus on the division rate fluctuations is in contrast to previous efforts to understand evolutionary dynamics in the presence of age-structure and non-genetic variability (see [20, 21]) which consider models with discrete age classes and assume the age distribution to be in steady state in order to obtain an effective SDE. We demonstrate that such approximations are not valid for physiologically realistic models of microbial growth. Therefore, it cannot be assumed that dynamics can be characterized by the constant growth rate terms obtained from the *steady-state phenotype* distribution and constant division rate; see Figure 1 (B) and (C).

The remainder of this paper is structured as follows. In section II, we start with a generalization of the Moran process derived from physiological models of growth, division, and cell-size control, and derive a Fokker-Plank equation for the genotype frequencies. Section III discusses Neutral dynamics in the Moran process. We will show that the rate of neutral genetic drift depends not only on the long-term fitness of a genotype, but also the variability in generation times, with larger variability leading to larger genetic drift. In section IV, we derive the fixation probabilities of a mutant arising in a large population, showing that the selection coefficient appearing in Eq. (3) is insufficient to predict the fate of a mutant. In particular, we will find that the fixation probability can decrease with the variability in generation times in the mutant population when long-term fitness is kept constant but is unaffected by that of the resident cells. In contrast, it is well known that variation in generation times tends to be beneficial from the perspective of long-term fitness (assuming constant mean generation time). Thus, there are situations where decreasing the selection coefficient can actually increase the fixation probability and vice-versa. As we will show, this can influence qualitative features of long-term evolutionary dynamics, leading to non-linear fitness trajectories in situations where classical theory predicts linear increase in fitness with time.

## II. CONTINUOUS AGE-STRUCTURED MORAN PROCESS

Consider a population consisting of *n* cells of genotype *m* (the mutants) and *N* − *n* cells of genotype *r* (the residents) growing according to the Moran process model, with the exception that instead of cells dividing at a constant probability per unit time, their divisions times are determined by a physiological model. To be precise, we assume that after living for a (possibly random) time *τ*_*d*_, called the *generation time*, a cell divides to produce two new offspring which themselves go on to produce offspring in the same manner. All the information about the underlying cellular physiology is encoded in how we select the generation time. For any microbe where the sizes of cells are controlled, the time at which a cell divides must be coupled to the cell’s volume; otherwise, small variations in doubling times result in unbounded fluctuations in size [3]. As a result, the model of generation time must account for cell sizes and growth rates, so the probability for a cell to divide in a given time interval after having survived to an age *a* depends on its size at birth *s*_*b*_, its growth rate *λ*, and its age *a*. We denote this probability by *γ*(*a, λ, s*_*b*_) *dt*. Letting *f* (*τ, λ, s*_*b*_) be the joint distribution of generation times, growth rates and initial cellsizes, we have 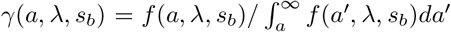. In Appendix A, we discuss a common framework for modeling cell-size regulation.

For each genotype *i*, we can define the per capita population division rate *ρ*_*i*_(*t*) at time *t* as the population average of single-cell division rates *γ*_*i*_:

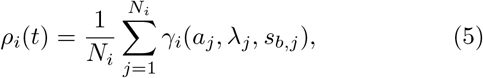

where the subscript *j* represents an individual cell in the population, and *N*_*i*_ is given by *n* for the mutant cells and *N* − *n* for the resident cells. Notice that only in the special case of exponentially distributed generation times (i.e. *f* (*τ, λ, s*_*b*_) = *λe*^−*τλ*^) do we obtain a time-independent per capita division rate *ρ*_*i*_ = *γ*_*i*_. However, it has previously been shown that in a deterministic treatment of the population dynamics which assumes selection is dominant over genetic drift (*ϕ ≫* 1*/*(*Ns*)) there is a well defined phenotype distribution,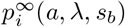. As it was shown in ref. [17], this steady-state distribution is the same as the steady-state distribution of single-cell variables in an exponentially growing population. This suggests it might be reasonable to neglect fluctuations in *ρ*_*i*_(*t*) and replace Eq. (5) with the average over this distribution:

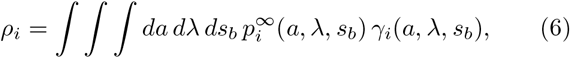

where now, *ρ*_*i*_s are time independent.

We consider the dynamics of the fraction of mutant cells, *ϕ*(*t*) ≡ *n/N* under the assumption that *ρ*_*i*_ are good approximations for the population division rates *ρ*_*i*_(*t*). The number fraction *ϕ* increases when the division of a mutant cell is followed by a resident cell getting discarded and decreases when the division of a resident cell is followed by a mutant cell getting discarded. Therefore, the probability *P* (*ϕ, t*) of observing *ϕ*(*t*) = *ϕ* at time *t* satisfies the master equation

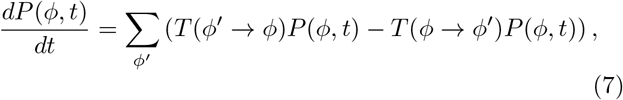

where the transition rate *T* (*ϕ* → *ϕ*^*′*^) from *ϕ* to *ϕ*^*′*^ can be written in terms of *ρ*_*i*_s as

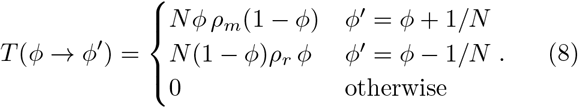

For large *N*, the Kramers–Moyal expansion of Eq. (7) leads to a Fokker-Plank equation whose corresponding stochastic differential equation is the continuous Moran process described in Eq. (1) (see Appendix B for derivation).

The continuous Moran process described in Eq. (1) is known to hold for a Markovian model of cell-division where cells divide with equal probability at any time independent of their age or size, but now, it is derived for an arbitrary model of growth, division, and cell-size regulation, assuming fluctuations in the division rates can be neglected.

In the next section we analyze the special case of neutral genetic drift, where 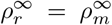, and show that simulation results disagree with the prediction from constant division rate assumption. As we will show, the rate of neutral genetic drift, as observed in simulations, is slower than what is predicted by the standard theory, even in the limit *N* → ∞. This disagreement reveals that, unexpectedly, the distribution of single-cell quantities never reach a steady state in constant-size populations. We will understand this insight in terms of coalescent theory as discussed in the next section.

## III. NEUTRAL EVOLUTION

We start by analyzing the model introduced in the previous section for the case of neutral mutations. Consider a population of *N* total cells of two genotypes of resident and mutant cells, *r* and *m*, both dividing with the same division rate *ρ*. From Eq. (1), the dynamics of the number fraction *ϕ* of the mutant species *m* in the classical Moran (that is, assuming generation times are exponentially distributed), is

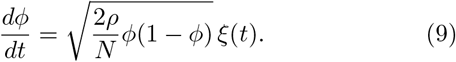

The dynamics of variance of *ϕ* in this case is given by Eq. (2) (see Appendix C1 for derivation); thus, the variance increases at a rate 2*ρ/N* (see Fig. 2). To compare this to the age-structured setting, we simulate a finite population of cells each growing exponentially in size with their elongation rates independently and identically distributed.

**FIG. 2.**
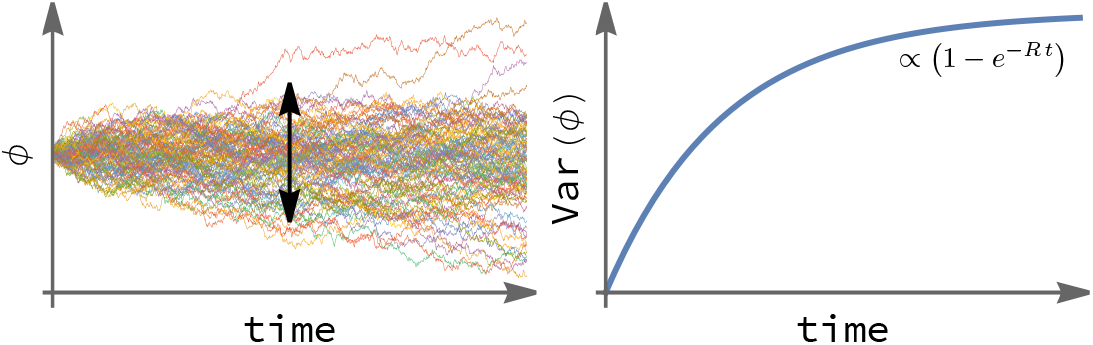
(Left) Stochastic trajectories of number fraction *ϕ* of a mutant starting from an initial number fraction *ϕ*_0_. In neutral genetic drift, the expected value of *ϕ* remains constant, but the variance increases over time. (Right) The exponential rate *R* with which the variance of *ϕ* increases over time is defined as the rate of neutral genetic drift.

We start with a simple model of division, where all cells are born with the same size *s*_*b*_ and divide with a fixed size *s*_*d*_ = 2*s*_*b*_. Assuming cells grow exponentially, *s*_*d*_ = *s*_*b*_*e*^*λτ*^ and hence *τ* = ln(2)*/λ*. Therefore, all the variation in the generation times comes from the growth rate *λ*, which we assumed to be uncorrelated between cells. For mathematical simplicity, we select an inverse-gamma distribution for *λ*, which results in a gamma distribution for *τ* with probability density

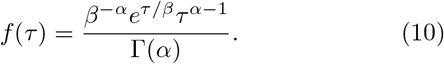

The Gamma distribution fits the empirical distribution of cell-size doubling times at least as well as a Gaussian distributions in various growth conditions, but also has many theoretical advantages: A gamma distribution has positive support, the Euler-Lotka equation (Eq. (4)) is analytically solvable for gamma distribution, and the gamma distribution naturally interpolates between exponential distribution (Markovian limit) and *δ* distribution (deterministic limit).

Simulations of the neutral model were performed using gamma distributed generation times for different coefficients of variations (CV is defined as the ratio of standard deviation to the mean). Figure 3 shows the ratio of the rate of neutral genetic drift, measured from these simulation, to the predicted value from Eq. (9). Here it can be seen that the variability in generation time increases the rate of genetic drift independently of how it affects the long-term fitness *ρ*, contradicting the analytical prediction.

**FIG. 3.**
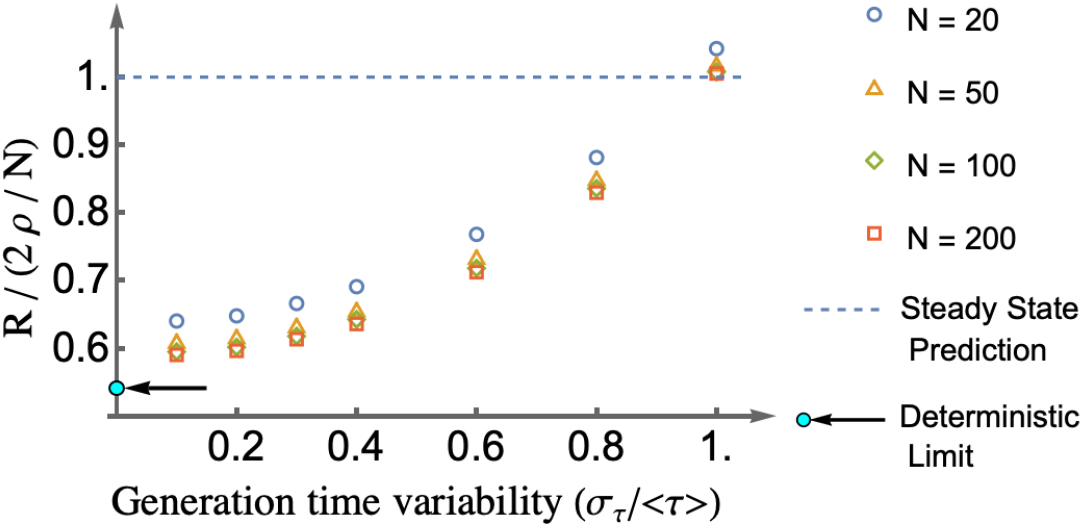
The ratio of the rate *R* of neutral genetic drift (as defined in Fig 2) from simulation to its predicted value 2*ρ/N* from the constant division rate assumption, Eq. (9), showing significant deviation from the predicted results. Different markers indicate different system sizes showing that the deviation is clearly not a finite-size effect. As the coefficient of variation of generation times approaches 1, the distribution of generation times approaches an exponential (Markovian limit) recovering the predicted results at large system sizes. For a smaller variability in generation times, fluctuations in the division rates have a significant effect on the rate of neutral genetic drift causing the deviation from the analytically predicted result.

At the Markovian limit *CV* = 1, where the generation time distribution is exponential, the simulated results approaches the prediction from Eq. (9) for large system sizes. For *CV <* 1, the fact that the rate of genetic drift at large system sizes approaches a different limit indicates that the division rate cannot be replaced by its average *ρ*.

### Persistent division rate fluctuations emerge from coalescence

To understand why division rate fluctuations are important, we consider a population consisting of only one species. While the number of cells is kept constant, the division rate and distribution of sizes and ages of the cells in the population can vary over time. Figure 4 shows the division rate for such a population, which appears to oscillate in populations of various sizes even after 1000 generation times. To ensure that these oscillations are not side effects of an oversimplified model, we have performed these simulation using a realistic model and parameter values for growth, division, and cell-size control of *E. coli* previously used in Refs [11, 12], although the same oscillations can be observed in the model with independent, gamma distributed generation times. They are reminiscent of the oscillations in growth rates of exponentially growing populations starting from a single cell [11] that are due to cells in the population sharing a common ancestor and dividing more or less at the same time. In growing populations however, oscillations decay over time, because as more generations pass since the cells share a common ancestor, the noise in the doubling time accumulates giving rise to less synchronized populations. Why does that not happen in populations with constant populations size? The key to answer this question lies in coalescent theory.

**FIG. 4.**
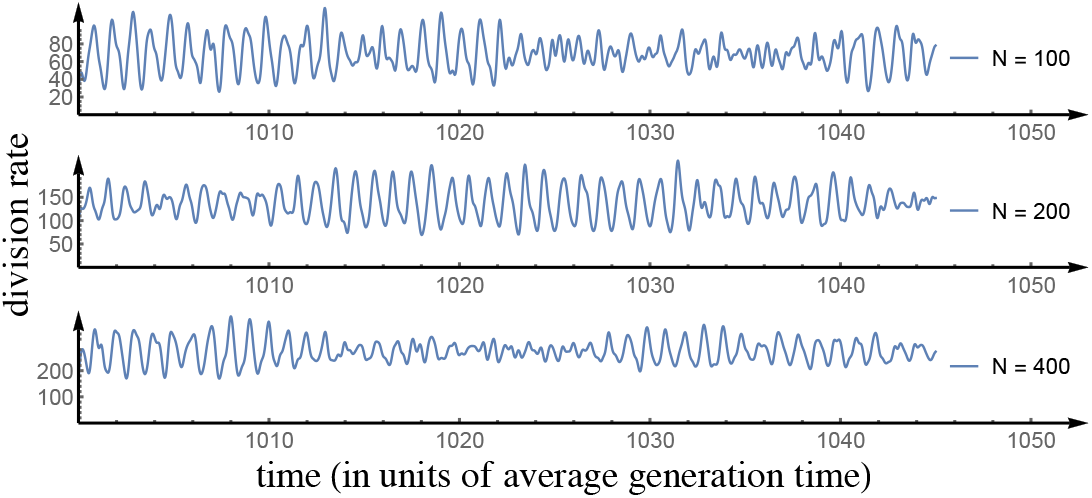
Stochastic spontaneous oscillations of division rates in simulations of constant-size populations of cells with a realistic model of growth, division, and cell-size regulation for *E. coli* from Ref. [11] for three different systems sizes *N* ∈ {100, 200, 400}, after 1000 generation from the start of the simulations. Simulation details: 10% division noise (size additive), 7% growth rate noise, adder model of cell-size regulation.

As shown in Fig. 5, starting from a population of cells with unknown history, after some time *t*_1_, the descendants of all cells except one will go extinct and the population will have a most recent common ancestor at some time *t*_1_ − *τ*_1_. Since these cells share a common ancestor, their division times are more or less synchronized. If we wait some additional time until time *t*_2_, one might expect that the population would be less synchronized because it has been more generations since the cells shared a common ancestor, but that is not true. As highlighted in the bottom panel of Fig. 5, the population at time *t*_2_ has a more recent common ancestor at the time *t*_2_ − *τ*_2_. Note that although the two so-called coalescence times *τ*_1_ and *τ*_2_ (times since the most recent common ancestor of the population at times *t*_1_ and *t*_2_) are stochastic and not necessarily the same, they have the same expected values, and therefore the population (on average) maintains the same degree of synchrony of cell-divisions over time. This synchrony gives rise to oscillations in the division rate and traveling waves in the cell-size and cell-age distributions of the population.

**FIG. 5.**
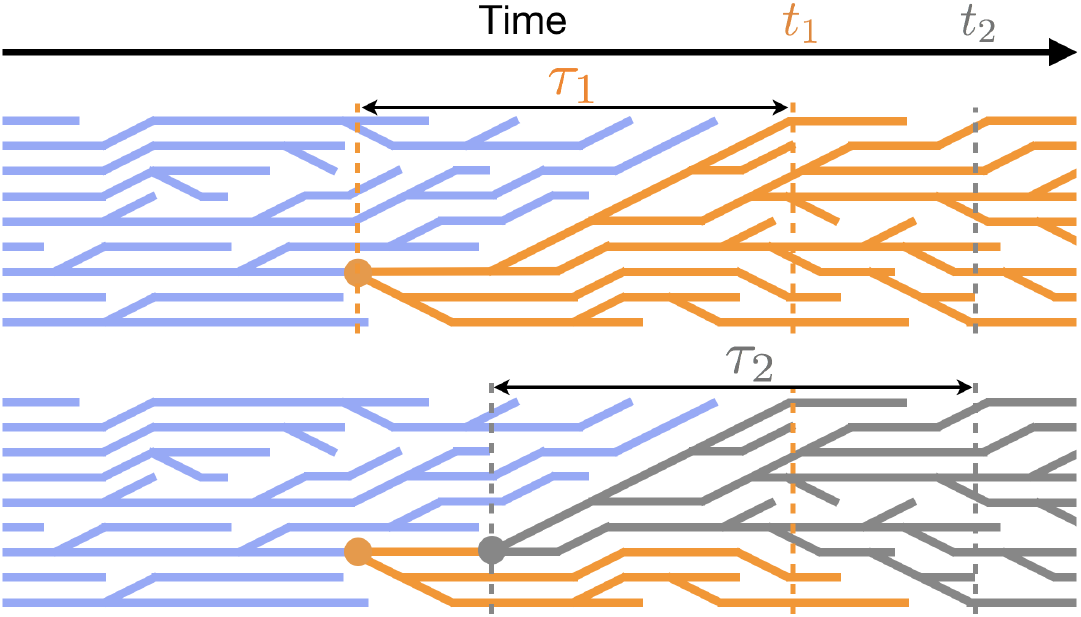
Top panel shows a sketch of the lineage history of a constant-size population. The population eventually reaches a state where all cells share a common ancestor (at time *t*_1_ in this illustration). The cells at this time share a most recent common ancestor from the time *t*_1_ − *τ*_1_, and therefore have some level of synchrony in their division times. One would expect that if we wait until some later time *t*_2_, cells would be less synchronized, as it has been more generations since they shared a common ancestor. However, as shown in the bottom panel, the cells at time *t*_2_ share a more recent common ancestor at time *t*_2_ − *τ*_2_. The two stochastic variables *τ*_1_ and *τ*_2_ are coalescence times for the population at times *t*_1_ and *t*_2_ and have the same expected values.

In the context of genetic drift, memory in the division rate prevent us from representing the effects of all single-cell variables by a single fitness variable. In particular, the variability in single-cell doubling times associated with a genotype affects the fate of an organism independently from how it affect the fitness variable *ρ*. In many bacteria such as *E. coli* this variability is much smaller than its mean, with the CV between 0.05 to 0.2 [22]. The rate of neutral genetic drift at these CV values is approximately the same as the zero-variability limit of Fig. 3. In this limit, the number fractions still experience a stochastic drift that can be approximated by (see Appendix C2 for the derivation)

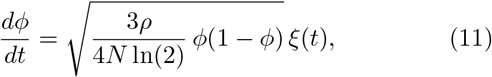

where *ρ* is the average division rate. The rate of neutral genetic drift in this case is *R* = 3*ρ/*4*N* ln(2) which is a factor 3*/*8 ln(2) ≈ 0.54 smaller than predicted by neglecting division rate fluctuations. This discrepancy can be traced back the fact that cells more or less preserve the order of their divisions at different generations when the variability in their elongation rates is very small.

## IV. ADAPTIVE EVOLUTION

In the previous section we found that the rate of genetic drift is determined not only by the long-term growth rates of species, but the details of the single-cell dynamics. We now shift our focus to adaptive evolution, starting with the question of how details of single-cell dynamics influence fixation probabilities. Consider a new mutant arising in a resident population described by the Moran process and let *ρ*_*r*_ and *ρ*_*m*_ be the long-term growth rates of the mutant and resident respectively. It will be assumed throughout that *ρ*_*m*_ *> ρ*_*r*_. If the mutant population survives to become sufficiently abundant, then its frequency *ϕ* will eventually be described by the deterministic logistic equation *dϕ/dt* = (*ρ*_*m*_ − *ρ*_*r*_)*ϕ*(1 − *ϕ*). Genetic drift plays a negligible role in this phase of the dynamics and fluctuations in the division rates can be ignored. However, when the mutant clone size is smaller than 1*/s*, the stochastic effects dominate and we need to consider how these may be influenced by details of the single-cell dynamics.

In order to obtain analytical results, we once again assume that the mutant population is described by the independent generation time model with a gamma distribution of division times, and consider the limits CV = 1 and CV ≈ 0. In the limit CV = 1, the mutant cells initially divide at a rate *ρ*_*m*_ so there is no distinction between the initial and long-term dynamics. In this case, the probability that the mutant fixes is obtained from the classical result of Haldane and Kimura shown in Eq. (3). Now consider CV ≈ 0. The long term growth rate of the mutant population is *ρ*_*m*_ = ln(2)/ ⟨*τ*⟩; however, initially the mutant cells divide approximately synchronously at a rate 1/ ⟨*τ*⟩. Since the convergence to the steady-state growth rate is extremely slow, we expect that the quantity *ρ*_*m*_*/* ln(2) = 1/ ⟨*τ*⟩ > *ρ*_*m*_ might replace *ρ* in the formulas for the fixation probability, leading a higher chance of fixation than the predictions from Kimura’s formula. This is indeed the case, in-fact, as we show in the next section the fixation probability of the synchronously dividing mutant populations is increased by a factor of 2 ln(2) compared to the prediction of Kimura’s formula. We will then generalize this to any value of CV and discuss the implications for long-term evolutionary dynamics.

### A. Derivation of fixation probabilities

We now derive a general formula for the fixation probability in terms of the generation time distribution. Assuming cells have independent generation times and the population is large enough that mutants are very unlikely to be expelled by the division of the other mutants, the probability that both of the cells offspring go extinct is simply 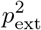. Therefore, letting *q* denote the probability that the mutant is expelled before it divides, we have

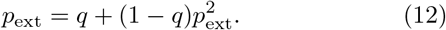

The extinction probability *p*_ext_ is the smaller root of the quadratic equation (approaching the fixed point from below) given by

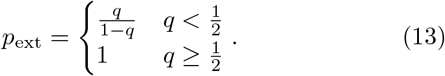

Letting *τ*_*d*_ be the doubling time of the mutant and *τ*_*e*_ the time until the mutant is expelled, *q* can be expressed as

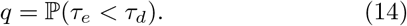

Since we are assuming the population is so large that nearly all removal events result from cells getting replaced by resident cells, the time until the mutant is expelled is simply division rate of resident cells, *ρ*_*r*_. In contrast to the mutant population, the division rate fluctuations of the resident population can safely be neglected, since they make only a higher order contribution to the expulsion of mutant cells. Hence *τ*_*e*_ is exponentially distributed with ⟨*τ*_*e*_⟩ = 1*/ρ*_*r*_. The distribution of *τ*_*d*_ is given by the generation time distribution, *f*_*m*_(*τ*). It follows that

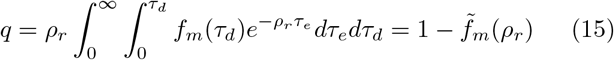

where 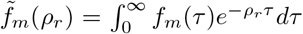 is the Laplace transform of *f*_*m*_ evaluated at *ρ*_*r*_. In order to make the connection to the Euler-Lotka equation, it is convenient to write Eq. (13) in terms of 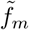 (see Fig. 6)

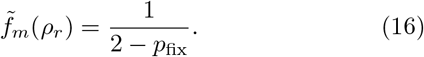

**FIG. 6.**
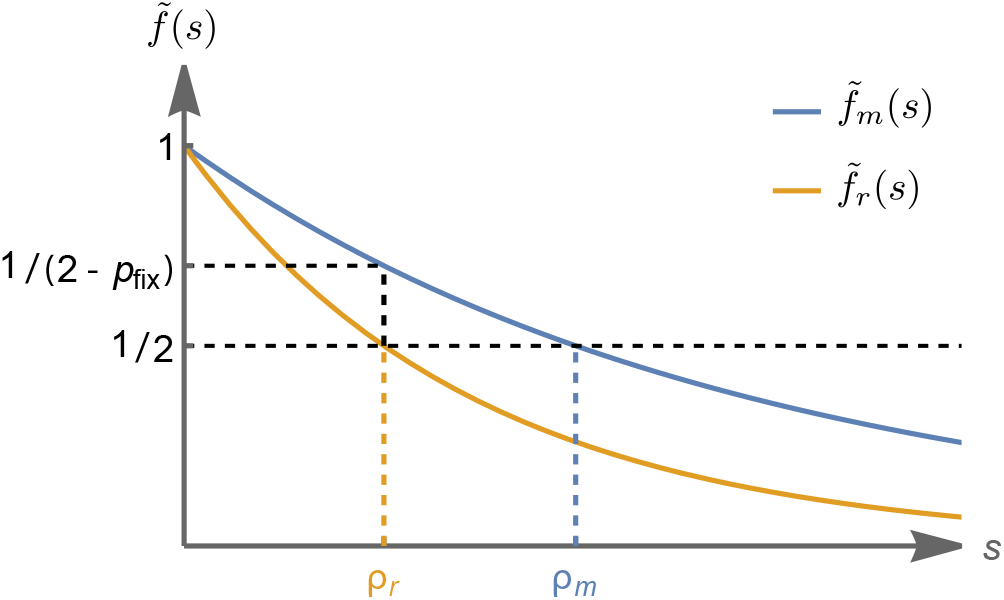
The relation between fixation probability and the Euler-Lotka equation: Euler-Lotka equation states that for both mutant and resident species *m* and *r*, the Laplace transforms of the generation-time density distributions 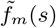 and 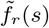 evaluated at their steady-state reproductive rates *ρ*_*m*_ and *ρ*_*r*_ are equal to 1*/*2. Equation (16) states that the Laplace transform of the generation-time density distribution of the mutant cells *f*_*m*_(*s*) evaluated at the reproductive rate of the resident cells *ρ*_*r*_ is given by 1*/*(2 − *p*_fix_).

Since 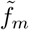 is equal to the right-hand side of the Euler-Lotka equation (Eq. (4)) with *ρ* replaced by 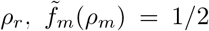 and 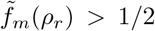 if and only if *ρ*_*r*_ *< ρ*_*m*_. Hence, as we would expect, *p*_fix_ *>* 0 (the mutant is beneficial) only when *ρ*_*m*_ *> ρ*_*r*_.

From this expression we can make three key observations: (1) As *N* → ∞ the fixation probabilities are coupled to the resident population’s growth dynamics only through the population growth rate, *ρ*_*r*_; however, (2) the fixation probabilities may depend in more complex ways on the mutant population’s growth dynamics because of the appearance of *f*_*m*_(*τ*). (3) Whether a mutant is deleterious or beneficial is determined by the long-term growth rates. That is, the fixation probability is positive if and only if *ρ*_*m*_ *> ρ*_*r*_.

This last statement can also be deduced from the following intuitive argument: Suppose *ρ*_*m*_ *> ρ*_*r*_. Let *ρ*_*m*_(*t*) be the transient division rate. We know there is some *n*_*c*_ such that when the mutant population size *m > n*_*c*_, *ρ*_*m*_(*t*) *> ρ*_*r*_ and at this point the population size *m* will increase approximately deterministically. There is some finite probability that *m* will exceed *n*_*c*_ even in an infinitely large population (since *n*_*c*_ can be reached through a finite sequence of cell divisions), thus there is a finite probability the mutants population will fix. Now suppose *ρ*_*m*_ *< ρ*_*r*_. In this case there is an *n*_*c*_ such that *ρ*_*m*_(*t*) *< ρ*_*r*_, thus by a similar argument the mutant will never fix. Since the mutant never fixes, it eventually go extinct with probability one. Notice that we have not used the independence of generation times in this argument, thus this result applies to more general models in which generation times are correlated.

### B. Markovian case

One case that has been well studied is when the distribution of generation times is exponentially distributed:

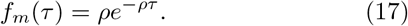

In this case the per unit time, per capita division rate is completely independent of the distribution of ages. Using Eq. (16), it is straightforward to derive the well-known result originally obtained by Haldane:

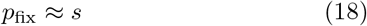

where *s* is the selection coefficient defined in Eq. (3).

### C. Deterministic case

We now consider the limiting case where the distribution of generation times is nearly a delta function around *τ*_*d*_. Thus, all cells in the mutant population divide at almost exactly the same age. In this case, Eq. (15) becomes

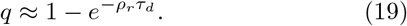

Here, we are neglecting any contribution from the variation around *τ*_*d*_. Using that the long-term growth rate of the mutant is *ρ*_*m*_ ≈ ln(2)*/τ*_*d*_, we obtain

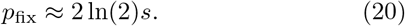

In this instance there is an additional factor of 2 ln(2) compared to the previous example. As we have suggested earlier, this discrepancy results from the fact that in the present case, the initial age distribution of the mutant population, and hence the division rate is distinct from that of the same genotype under stable exponentially growth. It is this transient division rate that determines the ultimate fate of the mutant population. It should be noted that the effect on fixation probabilities due to out of steady state dynamics is small and many other extrinsic factors can contribute to the prefactor of *s* in the fixation probability. However, a key point here is that the details of single-cell dynamics influence fixation probabilities and this has implications for long-term evolution dynamics, which we will discuss after generalizing the results above.

### D. General results for Gamma distributed division times

The examples above can be seen as special cases of a more general result obtained assuming a Gamma distribution of generation times. Then Eq. (15) becomes

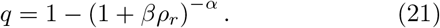

Using Eq. (4), we find that

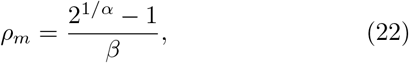

and *q* can be rewritten as

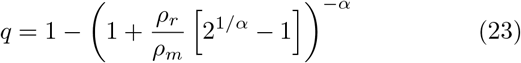

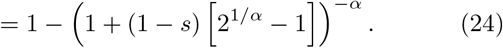

Using that 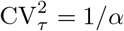, we can express *p*_fix_ solely in terms of the population growth rates and the coefficient of variation of mutant generation times:

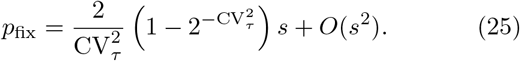

For 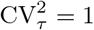 we obtain the Markovain case, *p*_fix_ = *s*, while in the limit 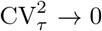 we retrieve the deterministic case *p*_fix_ = 2 ln(2) *s*. Agreement with numerical simulations is shown in Fig. 7.

**FIG. 7.**
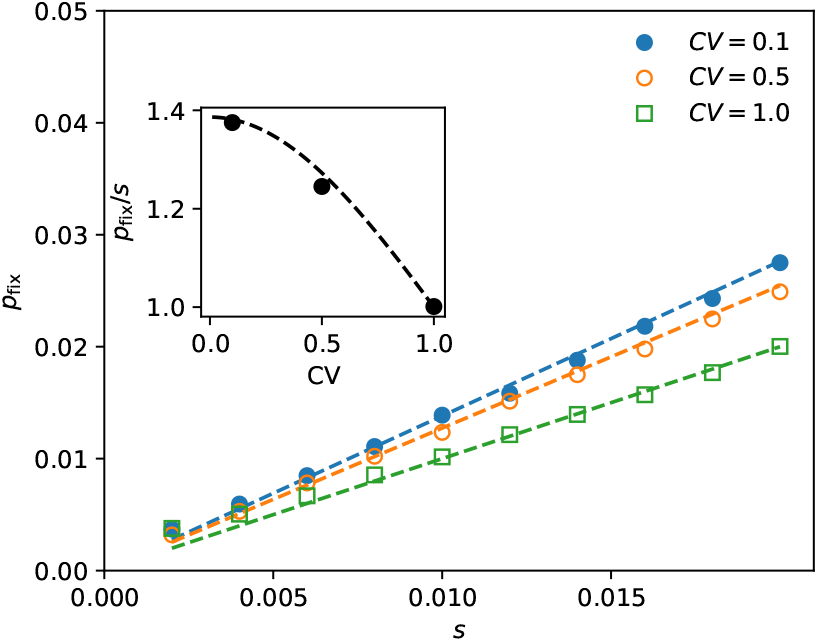
Numerical simulations of the fixation probability. Populations starting with one cell were run for 10^8^ runs and the fixation probability was computed from the fraction of populations that reaches a critical size where genetic drift ceased to dominate the dynamics. Dashed lines show the theoretical prediction of Eq. (25).

### E. Implications for long-term evolution

In the previous section we considered the dynamics following the emergence of a single mutation. We now consider evolutionary dynamics over long periods of time in which multiple mutations accumulate. To this end, we introduce the mutation probability *μ*, defined as the probability per cell-division for a mutation to occur. We assume, for simplicity, that each mutation increases the growth rate by an amount *δρ*. If a mutation does eventually fixate, it will take a time *τ*_sw_ on the order of 1*/δρ* ln *Ns* to sweep through the population [23]. We assume *τ*_sw_ ≪ 1*/*(*μNρs*), or *μN* ln *Ns ≪* 1 [24]. This regime is often referred to as the strong selection-weak mutation regime and it ensures mutations are unlikely to complete with each other and can therefore be treated as independent events.

A basic question one can ask regarding the evolutionary trajectories is: How will the growth rate increase over time in such a population? In particular, one typically considers the dynamics of the growth rate averaged over many realizations of an evolving population, which we denote ⟨*ρ*⟩. The per unit time increase in ⟨*ρ*⟩ is simply the rate at which mutations emerge multiplied by the probability of fixation; that is,

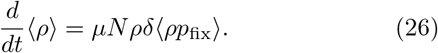

Using the classical formula *p*_fix_ = *s*, we find *d ⟨ρ⟩ /dt* = *μN* (*δρ*)^2^ and hence under these assumptions the long-term growth of ⟨*ρ*⟩ is linear.

As we have shown, the fixation probability will depend not only on *ρ* but the details of single-cell growth and division. As a contrived, but illustrative example, suppose each mutation increases 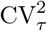 by a fixed increment 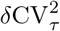 and the generation time by an increment *δτ* which is selected to ensure *δρ* remains constant. As the population adapts, the CV increases. This decreases the fixation probabilities and as a result adaptation slows down. In Figure 8 we show simulations of this model which illustrate this point. In reality the growth rates in an evolving population will not increase linearly for many other reasons, such as epistasis. However, our results suggest that the transient dynamics of mutant populations are also a factor which influence the long-term evolutionary dynamics. We leave it to future work to explore this effect within the context of more realistic models.

**FIG. 8.**
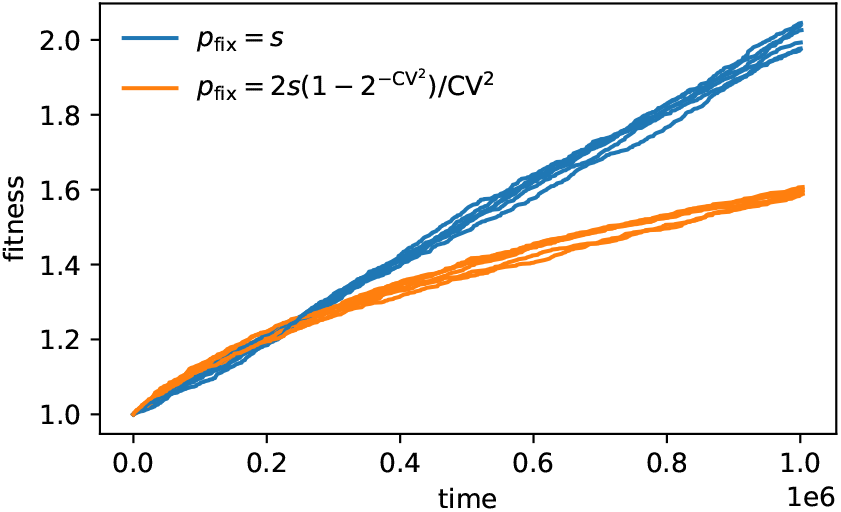
Numerical simulations of the long-term evolutionary dynamics using Kimura’s formula (blue line) and the corrected formula which accounts for division rate fluctuations (orange). Each line is a separate simulation. When accounting for the effects of fluctuations on fixation probabilities, evolution slows down considerably. This is because the coefficient of variation increases as the population adapts, decreasing the chance for new mutants to fix. In our simulations, the times between mutations is drawn from an exponential distribution with rate parameters *μNδρ*^2^*p*_fix_. Parameter values used are *s* = 0.01 *μ* = 10^−6^, *N* = 10^3^, ensuring we are in the strong selection-weak mutation regime.

## V. DISCUSSION

With recent advances in single-cell technology [22, 25] have come very detailed models of single-cell growth, division, and size regulation [26]. Population dynamics studies based on these models [10–13, 17, 27–30] have determined which details of these models affect population level quantities. In particular, we have learned that the details of single-cell elongation dynamics (but not details of division process) affect the growth rate of populations [11–13]. Recent work has drawn attention to the persistence of out-of-steady-state dynamics of a population of cells starting from a single cell [11, 31, 32]. These out-of-steady-state dynamics manifest in forms of oscillations in the population growth rate and traveling waves in distributions of sizes and ages of the cells, and they exist because descendants of a cell maintain a degree of synchrony in their growth and division. Their persistence time in exponentially growing populations also only depends on single-cell elongation dynamics and not the details of cell-division and cell-size regulation [11]. Here, we have shown that for finite populations, similar fluctuations in the division rate persist infinitely long (also observed in Ref. [33]), and they decrease the rate of genetic drift of neutral mutants. We leave it to future work to develop an analytical understanding of how the generation time distribution influences the rate of genetic drift in the Moran model.

In the context of beneficial mutants, we have shown that variability in doubling time reduces the probability of fixation, but it is always larger than the classical results from the Moran process. The analytic expression obtained for fixation probability has an interesting relationship to the Euler-Lotka equation as shown in Fig. 6. The Euler-Lotka equation states that the Laplace transform of the doubling-time distribution evaluated at the population growth rate *ρ*_*m*_ is 1*/*2, while in our case, the Laplace transform of the doubling-time distribution evaluated at the population growth rate of the *resident* cells *ρ*_*r*_ is 1*/*(2 − *p*_fix_). Finally, we have shown that the small correction to the fixation probability due to out-of-steady state effects can have potentially large effects on the long-term evolutionary trajectories. For example, if populations adapt by increasing the variability in their generation times, the rate of adaptation will decrease.

## VI. ACKNOWLEDGEMENT

We thank Oren Raz, David Lacoste and Simone Pigolotti for helpful comments on this manuscript. This work was initiated at the Aspen Center for Physics, which is supported by National Science Foundation grant PHY-1607611. EL was supported by NSF grant DMS-1902895.

## Appendix A: Cell-size control model

We assume cells attempt to divide when they reach a target division size *f* (*s*_*b*_), which in general is a function of their initial size *s*_*b*_. The actual division size *s*_*d*_ is the target division size plus a noise term, *s*_*d*_ = *f* (*s*_*b*_) + *ξ*. We assume cells grow exponentially with a possibly random elongation rate *λ* such that the generation time *τ*_*d*_ satisfies 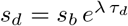.

Commonly considered candidates for the function *f* are *f* (*s*_*b*_) = 2*s*_*b*_ (the timer model), *f* (*s*_*b*_) = *s*_*b*_ + Δ (the adder model), 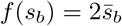 (the sizer model). The functional form of *f* is only relevant when the variance of the division noise *ξ* is nonzero; when this variance is zero, all three of these models are equivalent. With the exception of the simulation shown in Fig. 4 (where the adder model of cell-size regulation is used), we have neglected the division noise *ξ*, and considered the case where the only source of variability in the generation time *τ*_*d*_ comes from the variability in *λ*.

We refer to Ref. [3] for an in-depth discussion of the cell-size control model and its implications for population growth.

## Appendix B: Derivation of the Fokker-Planck equation for the number fraction of mutant species

Here we derive a Fokker-Plank equation by Kramers–Moyal expansion of Eq. (7) whose corresponding SDE is given in Eq. (1). We start by plugging the transition rates from Eq. (8) into the master equation Eq. (7). Defining *ϵ* ≡ 1*/N*, we have:

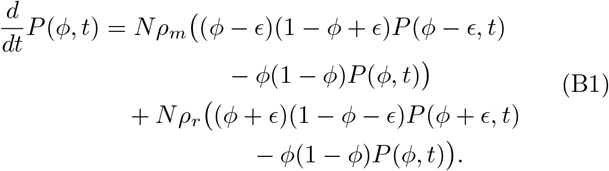

We can recognize this master equation as a difference equation by defining *f* (*ϕ, t*) ≡ *ϕ*(1 − *ϕ*)*P* (*ϕ, t*):

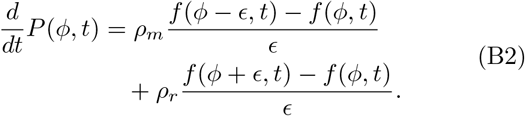

In the continuum limit, we can expand *f* (*ϕ* ± *ϵ, t*) up to second order in *ϵ* to derive the Fokker-Planck equation for the dynamics of the probability density *P* (*ϕ, t*) of the number fraction *ϕ* in time:

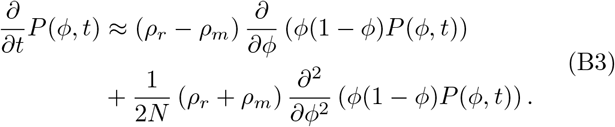

This equation describes the time evolution of the probability density of *ϕ* obeying the SDE given in Eq. (1).

## Appendix C: Rate of neutral genetic drift

We discussed in Section III that the simulation results deviate from the steady-state approximation for the rate of neutral genetic drift (which we derive below) except when the coefficient of variation (CV) approaches one, which corresponds to exponentially distributed doubling time or the Markovian limit. At the opposite limit, CV → 0, cells are expected to perfectly synchronize. Below, we derive the rate of neutral genetic drift at these two limits.

### 1. Steady state approximation

At the steady state, where the division rates *ρ*_*r*_ and *ρ*_*m*_ are time independent, we can use the Fokker-Planck equation given in Eq. (B3) to derive a differential equation for the rate of change of the variance of *ϕ* starting from some initial value 0 *< ϕ*_0_ *<* 1. In the neutral case *ρ*_*r*_ = *ρ*_*m*_ = *ρ*, we have

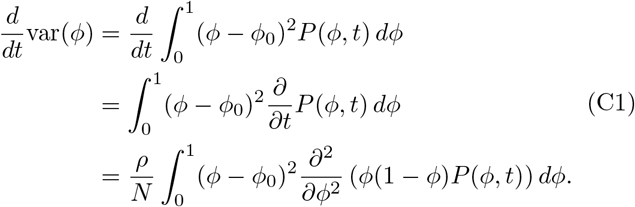

After integrating by parts twice, we have

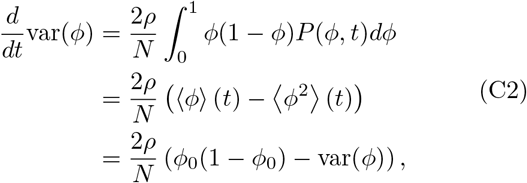

where we have used ⟨*ϕ*⟩ (*t*) = *ϕ*_0_. Solving for var(*ϕ*), we have

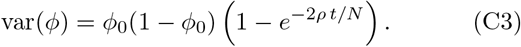

With the rate of neutral genetic drift given by *R* = 2*ρ/N*.

### 2. Deterministic limit

At the limit where the Coefficient of Variation (CV) of the cell-size doubling time goes to zero, we expect all the division events of each generation to happen at the same time. One might think that in this limit, the age-structured Moron process would approach the Wright-Fisher model where in discrete time intervals, all cells divide and the half of the population is discarded. That is not the case here. In the age-structured Moran process, cells divide one at a time and after each division, one cell is removed, and as a result, not all cells get to divide. This distinction survives even at CV → 0 limit where all the division events have to take place at the same time. To predict the rate of neutral genetic drift in this limit, we imagine the population of *n* mutant cells and *N* − *n* resident cells at generation *g* (note that in this limit, division times are discrete, and we can use generation numbers instead of time). In the next generation, we randomly choose previous generation cells to divide and discard cells to keep the population constant. We continue until no cells of generation *g* are present. Our goal is to find the expected value and the variance of number *n*^*′*^ of mutant cells in the next generation *g* + 1. This is a purely combinatorics problem. Below, we will show that the expected value of *n*^*′*^ given *n* is give by ⟨*n*^*′*^ |*n*⟩ = *n* using a symmetry argument. Also, for large *N*, we will show that the variance of *n*^*′*^ given *n* is

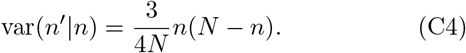

Given the conditional mean and variance of *n*^*′*^, at the continuum limit, we can approximate this processes as the following stochastic differential equation

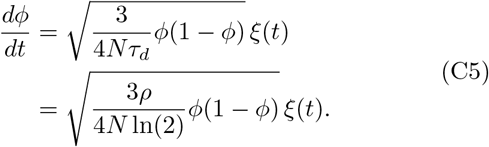

The rate of neutral genetic drift for this case is given by

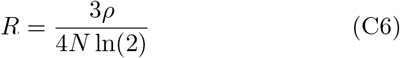

which is a factor 3*/*8 ln(2) ≈ 0.54 smaller than the rate of neutral genetic drift in the Markovian case *R* = 2*ρ/N*.

Note that the continuous time approximation made here does not become exact at the large *N* limit, given that the divisions clearly happen in discrete time intervals. Nevertheless, the estimated ratio of the rate of genetic drift matches the simulation fairly well as seen in the CV → 0 limit of Fig. 3.

#### Derivations of the mean and the variance of n^′^ given n

We use the symmetry that in the neutral case, all the cells are identical, and therefore, the numbers *X*_*i*_ ∈ {0, 1, 2} of their surviving progeny at each generation are identically distributed (albeit dependent) random variables. Since the total population is constant, we have 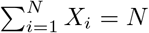. Therefore, 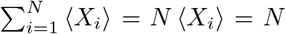 or ⟨*X*_*i*_⟩ = 1 for each cell *i*. The number *n*^*′*^ of the mutant cells in the new generation is given by sum of the numbers of the progenies of the previous generation mutant cells, 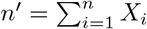, and therefore we have ⟨*n*^*′*^⟩ = *n*, showing that the expected value of the number of mutant cells stays constant throughout the neutral dynamics.

Finding the variance of *n*^*′*^ is a bit more tricky. We first show that the variance of the number of progenies of each cell Var (*X*_*i*_) is 3*/*4. But since *X*_*i*_s are dependent, we also need to find the Cov (*X*_*i*_, *X*_*j*_) (for *i* ≠ *j*) to find the variance of 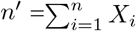. We will show that it is given by Cov (*X*_*i*_, *X*_*j*_) = −Var (*X*_*i*_)*/*(*N* −1). Finally, we will express the variance of *n*^*′*^ in terms of the variance and covariance of the variables *X*_*i*_ as

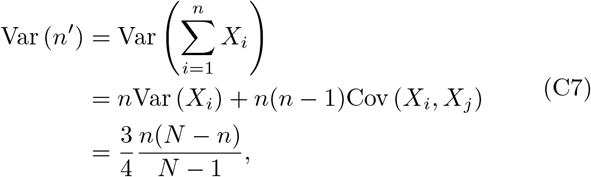

recovering Eq. (C4) in the large *N* limit.

The variance of *X*_*i*_ is given by

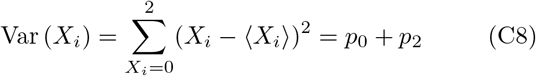

where *p*_0_ = *Prob*(*X*_*i*_ = 0) and *p*_2_ = *Prob*(*X*_*i*_ = 2). Since 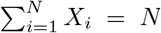, *p*_0_ = *p*_2_; that is for every cell with zero progeny, there is exactly one cell with two progenies. So let us find *p*_2_.

At each step of the process, a cell divides and another is discarded. We enumerate these steps with a discrete time variable *t*. We define the variable *f* to be the expected value of the fraction of generation *g* cells at time *t*. At each step, *f* decreases by 1*/N* due to division. Additionally, *f* decreases by 1*/N* if the discarded cell is from the previous generation (with the probability *f*). This gives the dynamics of *f* as

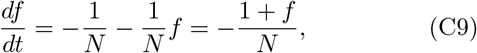

with solution *f* = 2*e*^−*t/N*^ − 1. This fraction goes to zero after *t* = *N* ln(2) steps. For a cell that divides when the fraction of the previous generation cells is *f*, the probability that it ends up with both of its progenies in the next generation is given by product of the probabilities 1 − 2*/N* of them not getting discarded in each step of division after *f*. The number of steps after fraction *f* is given by *N* ln(2) − *N* ln(2*/*(*f* + 1)) = *N* ln(1 + *f*). This gives the conditional probability of a cell ending with two surviving daughter cells given that it divided at *f* as

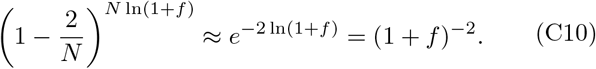

Since each cell of generation *g* eventually either divides or gets replaced as *f* decreases from 1 to 0, for each cell, *f* at which it divides or gets replaced is uniformly distributed. Conditioned on the value of *f*, the probability of division is (1*/f N*)*/*(1*/f N* + 1*/N*) = 1*/*(1 + *f*) (at *f*, the probability of division of each one of the remaining *f N* cells is 1*/f N* and the probability of them being discarded is approximately 1*/N*). Putting it all together, the probability *p*_2_ that a given cell ends up with two progenies is the integral over *f* of the probability that it divides at *f* multiplied by the conditional probability that neither of the daughter cells gets discarded given that it divided at *f*

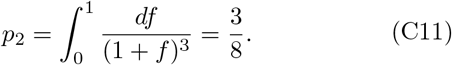

This gives Var (*X*_*i*_) = *p*_0_ + *p*_2_ = 2*p*_2_ = 3*/*4.

Next, we find the Cov (*X*_*i*_, *X*_*j*_) for *i* ≠ *j*. Let us define *δX*_*i*_ = *X*_*i*_ − ⟨*X*_*i*_⟩. The covariance can be written as

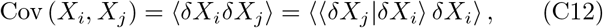

where ⟨*δX*_*j*_|*δX*_*i*_⟩ is the conditional expected value of *δX*_*j*_ given *δX*_*i*_. Since 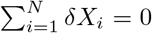, by symmetry, ⟨*δX*_*j*_|*δX*_*i*_⟩= −1*/*(*N* − 1)*δX*_*i*_. This gives

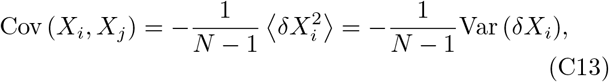

which is the last ingredient needed in Eq. (C7).

### 3. Code availability

Code related to this work can be found at [34].

